# High-Speed Automatic Characterization of Rare Events in Flow Cytometric Data

**DOI:** 10.1101/591644

**Authors:** Yuan Qi, Youhan Fang, David R. Sinclair, Shangqin Guo, Meritxell Alberich-Jorda, Jun Lu, Daniel G. Tenen, Michael G. Kharas, Saumyadipta Pyne

## Abstract

A new computational framework for FLow cytometric Analysis of Rare Events (FLARE) has been developed specifically for fast and automatic identification of rare cell populations in very large samples generated by platforms like multi-parametric flow cytometry. Using a hierarchical Bayesian model and information-sharing via parallel computation, FLARE rapidly explores the high-dimensional marker-space to detect highly rare populations that are consistent across multiple samples. Further it can focus within specified regions of interest in marker-space to detect subpopulations with desired precision.

## Introduction

Studies focusing on rare cell populations are becoming increasingly common owing to technological advances such as high-speed, multi-parametric flow cytometry, and emerging biomedical applications like stem cell therapy, and single cell analysis. Researchers in fields such as hematology, cancer, immunology, pathology, stem cell biology, and regenerative medicine, have focused on many interesting, yet relatively rare, populations of cells in blood and other tissues and systems that have important biomedical functions and characteristics, e.g., long-term hematopoietic stem cells.

Methods for accurate detection or automated isolation of rare therapy-resistant cells in tumors with stem cell like properties, tumor cells circulating in blood, or regulatory T cells, can have profound influence on basic and clinical research. Platforms like multi-color flow cytometry, in conjunction with the development of diverse panels of markers and antibodies, have been used to establish signatures for various rare cellular species and lineages in terms of the expressed surface and intracellular marker proteins [1]. Advances in mass cytometry have promised the ability to determine 50-100 features per cell [2,3]. To address such increasingly multi-parametric and multiplexed immunoprofiling of each cell, studies have demonstrated the critical need for systematic and automated multivariate analysis and visualization suitable for high-dimensional data [4–6]. As the number of potential combinations of markers continues to grow exponentially (with the number of markers), a thorough search for rare events in high-dimensional marker-space clearly gets difficult with the more subjective and painstaking approach of traditional manual gating [7].

Analytically, a population of cells having similar, characteristic expression of *k* (> 1) markers can be measured as events with similar fluorescence intensities, i.e., as a cluster of points located closely in *k*-dimensional marker-space [4]. However traditional clustering approaches may not be adequate for identification of rare cell populations for several technical reasons. The new data are not only high-dimensional (i.e., involving multi-parametric or multiplexed panels) but simultaneously are also high-resolution (single cell level) and considerably high-throughput (hundreds of thousands of cells per sample) by design. Typically, therefore, if a population of interest is rare and consists of, say, fewer than 1% or 0.1% of the total number of cells in a given sample, then for reliable detection of such a population, it is common to use a sample size (*N*) in the order of 10^5^ − 10^6^ cells, each measured as a *k*-dimensional point. Thus a large cytometric sample can present a “searching for a needle in a haystack” scenario for the identification of any rare population therein, resulting either in inefficient coverage of the *k*-dimensional marker-space (the volume of which increasing exponentially with *k*), or detection of a number of spurious small populations (often outliers of larger, noisy populations). In general, clustering methods like k-means or hierarchical clustering use some measure of distance between every pair of points to determine their closeness for clustering assignment. While effective for clustering a few thousands genes or features in omic data, clearly such quadratic-time *O*(*N*^2^) approaches would be computationally inadequate for searching complex cytometric datasets with much larger *N*.

Another practical challenge stems from biological and/or technological sources of inter-sample variation including single cell level heterogeneity, individual subjects, different time-points and conditions, and platform noise — all of which make consistent identification of particularly the rarer populations difficult. Moreover, as cells undergo state transitions, for instance during differentiation, the corresponding changes in marker-expressions result in hierarchies of inter-connected clusters. Such clusters may contain complex high-dimensional structures such as heavy tails or skewness, that present unique data modeling challenges for computational analysis [5,8]. Therefore, we developed FLARE as a new computational framework that can simultaneously meet the somewhat conflicting requirements of (a) high speed, (b) high precision, and (c) robust data modeling.

## Model

In this section, we describe the our new hierarchical Bayesian model, FLARE, for FLow cytometric Analysis of Rare Events, to identify cell populations from multiple samples and detecting rare cell populations. Given the increasing high-dimensionality of cytometric data, there is a critical need to assist the manual gating procedure using unsupervised computational approaches to explore the marker-space, especially to identify specific cell populations that may appear at unknown locations under certain conditions such as drug-resistant cells or a rare signature of disease prognosis.

To this end, we designed a hierarchical Bayesian model that can share information across multiple samples to substantiate the occurrence of any genuine rare cluster of events. First, we model the cell populations in each sample by a mixture of probability distributions, say, multivariate Gaussian components, so that we can assign a probability score to associate each cell with a population, thus reflecting the underlying structures of individual samples. Second, we let the Gaussian components — corresponding to cell populations in different samples — be similar to each other via common prototype populations up to certain small variations, so that we can capture the minor differences between individual samples. Third, we allow some Gaussian components to appear only in certain — but not necessarily all — samples, and report these populations, even if they are rare events.

Let us denote the cytometric data by **X** and the cell memberships by 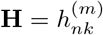 where 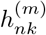 denotes the membership of the n-th cell in the m-th sample to that sample’s k-th Gaussian component with mean 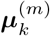 and precision 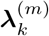. Then the data likelihood is

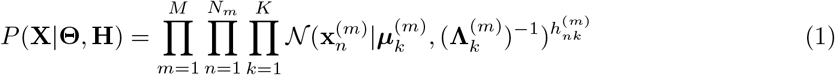

where *M* is the number of samples, *N_m_* the number of cells in sample *m, K* the maximal number of cell populations for each sample, and 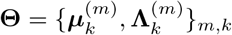. Each data point 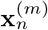 has dimension *D*. The latent membership indicators **H** has a factorized discrete prior distribution:

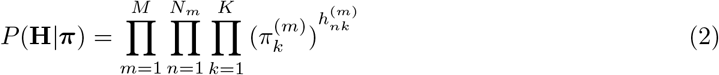

where 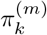 is the probability of the *k*th population appearing in sample *m* and 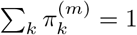. If 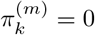, then the *k*th population does not exist in the m-th sample. To model the uncertainty in 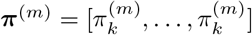, we use a symmetric Dirichlet prior distribution:

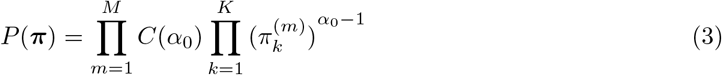

where *α*_0_ is a hyperparameter and 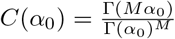.

To share information between clusters of different samples, we let the mean parameter, 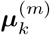, of each cluster in a sample follow a Gaussian prior distribution common to all samples:

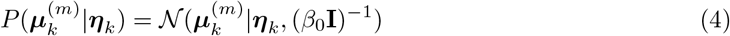

where ***η**_k_* is the mean parameter of the *k*th prototype cluster — which is estimated from data as 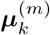 —and *β*_0_ is a hyperparameter. Similar, the covariance matrix, 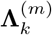, of each cluster in a sample follows a Wishart prior distribution common to all samples:

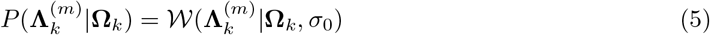

where **Ω**_*k*_ is a symmetric, positive definite matrix — estimated from data just as 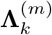 — and *σ*_0_ is the degree of freedom.

Since we need to estimate the parameters of the prototype clusters from data as well, we assign a Gaussian hyper-prior distribution over the mean of each prototype cluster, ***η_k_***:

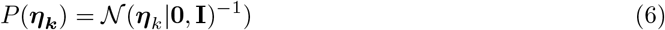

Also, we assign an Inverse-Wishart hyper-prior distribution over the shape of each each prototype cluster, **Ω**:

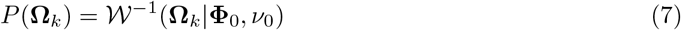

where **Φ**_0_ and *ν*_0_ are hyperparameters. In our experiments, we set **Φ**_0_ = **I** and *ν*_0_ = 6*D* to obtain a diffuse prior over **Ω**_*k*_.

Combining the data likelihood, the priors and the hyper-priors, we obtain the following joint distribution for our model:

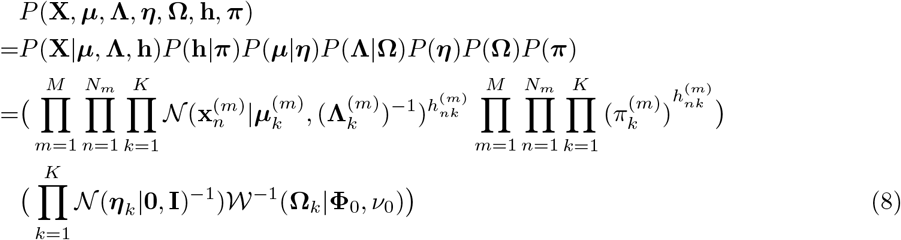

The joint distribution is depicted in Fig. 1.

**Fig 1.**
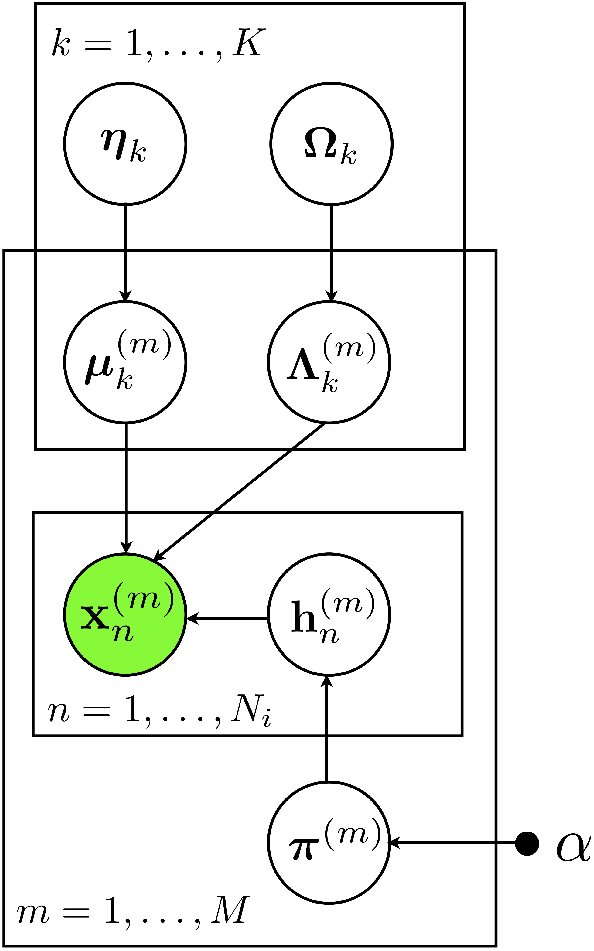
The probabilistic graphical model representation of FLARE. *x* is the cell, *μ* and Λ are the mean and covariance of the parent clusters, *η* and Ω are the mean and covariance of cluster *k, π* is the membership distribution, *h* is the membership indicator and *α* is the hyperparameter for *π*. The superscript *m* indicates the *m*^th^ sample.

With the priors specified over the cluster parameters in multiple samples and the hyper-priors over the parameters of the prototype clusters, we constructed a hierarchical Bayesian model, FLARE. The model allows the cluster locations (given by the means) and the shapes (given by the covariance matrices) of each sample to be similar to those of their prototype cluster so that the information from multiple samples could be combined for accurate and robust estimation of clusters. At the same time, FLARE allows the clusters of each sample to be slightly different from their prototypes, accounting for the variations among different biological samples. In our experiments, we set *β*_0_ = 500 and *σ*_0_ = 6*D* so that the stochastic variation between a sample cluster mean and its prototype cluster mean is reasonable small.

Notably, in our model, a cluster can also not contain any data point in a particular sample, and thus, the cluster may be absent in certain samples. From our estimation results, we can easily distinguish which clusters are common to all samples and which only appear in certain samples. The Variational Bayes (VB) approach to compute the approximate posterior distribution of the FLARE parameters is described in Section A of the Supplementary Materials. The algorithm for parallel computing, and specifically for for sharing information across samples to detect the consistent clusters is described in Section B of the Supplementary Materials.

## Results and discussion

We developed a new computational framework FLARE for FLow cytometric Analysis of Rare Events, although it may be applicable to other platforms that generate multi-marker data per cell.

FLARE is based on a hierarchical Bayesian model, and employs parallel computation for its high-speed high-precision analysis. The Bayesian model (Fig 2) of FLARE allows implementation of several distinct features to specifically address the challenges mentioned above. For consistent identification of a particular rare population *C*, the model parameters allow information about *C* to be shared across different samples. In our parallel computing framework, we implemented this via communication among nodes each of which analyzed a distinct sample. The strategy builds repeated inter-sample consensus on the existence of *C* (or lack thereof), thus guarding against unsupervised detection of possibly numerous spurious small populations. Consequently, the model estimation is robust against high inter-sample variation and platform noise, which otherwise are known to affect the reproducibility or the quality of match between analogous populations across samples and replicates [9].

**Fig 2.**
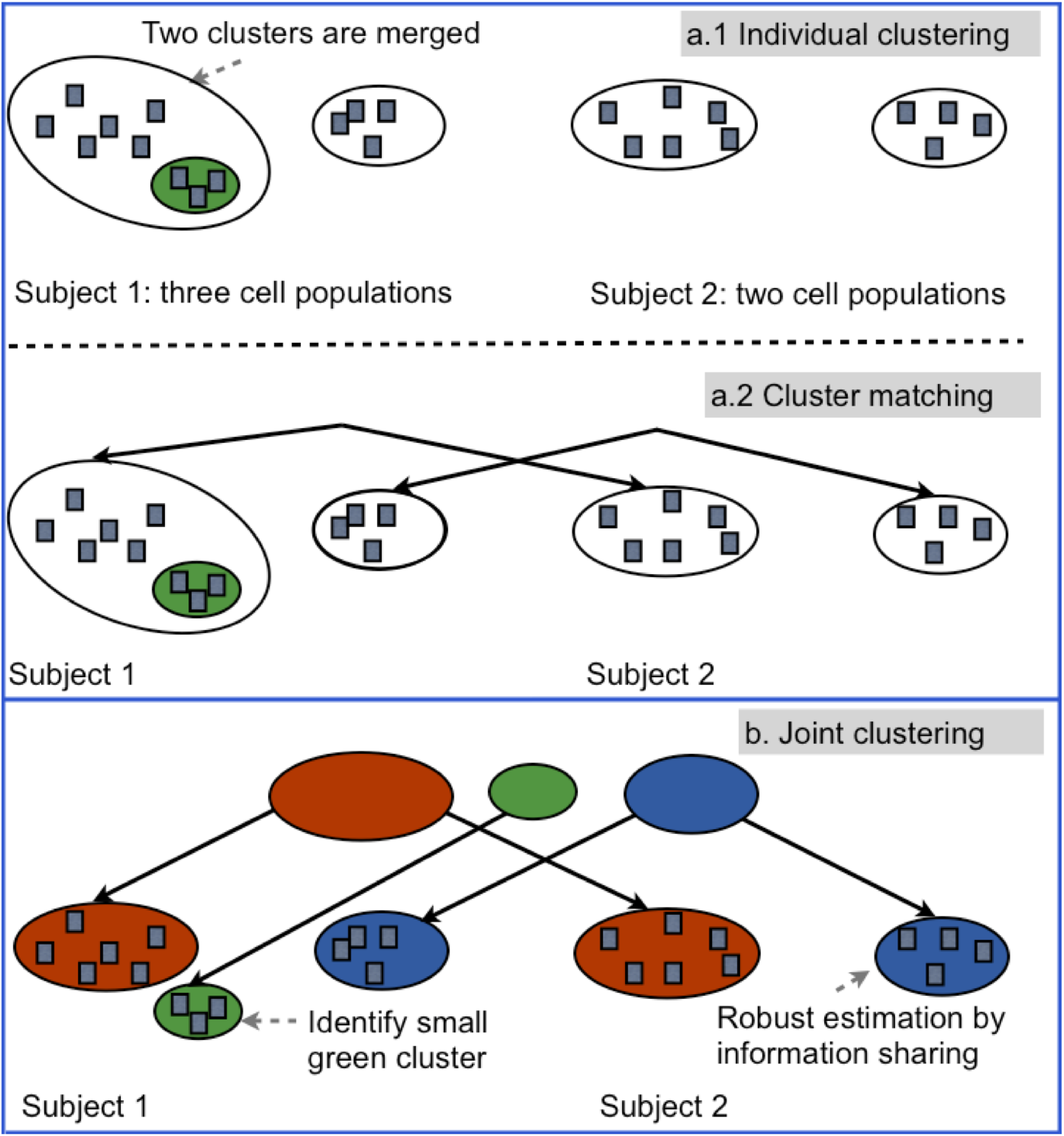
Illustrative example for FLARE and its graphical model representation. Panels a.1 and a.2 show the limitation of separate population analysis on individual samples: it misses the detection of the rare cell population in green. Panel b shows that by sharing population information via the parental nodes, FLARE can more accurately estimate the big cell population in the red cluster and also detect the rare population in the green cluster. Panel c describes the hierarchical Bayesian model of FLARE (see Section A of the Supplementary Materials for details).

We found FLARE’s information-sharing feature to be especially useful for rare cell populations (e.g., blue clusters in Fig 2) which contain very few cells since it effectively pools together more observations for estimation. Second, the estimation ambiguity (between the red and the green clusters in Sample 1, corresponding to Subject 1, in Fig 2) is reduced since the information about a population (e.g., the red cluster of Sample 2, corresponding to Subject 2, in Fig 2) can guide the estimation of its counterpart across samples (i.e., the red cluster of Sample 1 in Fig 2) in FLARE’s joint model. Third, the joint model also allows partial consistency such that some clusters can exist in one or more samples but not necessarily in all of them. Thus, without needing any additional cluster alignment [4, 13], we can identify also those clusters (e.g., the green cluster in Fig 2b) that exist only in certain samples, a situation that is not uncommon for rare populations e.g., transient subsets that are present only during certain stages of cell differentiation and are absent otherwise).

For our first application of FLARE, we generated a 6-marker cytometric dataset to study cells from mouse bone-marrow (Supplementary Section C). In the first step, without any human guidance, unsupervised analysis by FLARE was run on 14 “training” samples, including multiple biological and technical replicates, and it identified a subset bearing a 6-marker signature of long-term murine hematopoietic stem cells (LT-HSCs) (Fig 3a-b). In the second step, using this signature location parameter, FLARE focused on the corresponding region in an entirely new and much larger “test” sample (containing more than a million cells measured with the same 6 markers) to detect a very rare population (containing 0.045% of the total number of cells in the sample) with a more precise LT-HSC signature. This finding is supported by earlier analyses using sequential two-dimensional gating, according to which LT-HSCs are known to be Lineage^-^Kit^+^Sca^+^CD34^-^CD48^-^CD150+. Using parallel computation (see Section B of the Supplementary), the two steps took less than 10 minutes to finish. The consistency of the detected subsets across all 14 training samples, the small size of the final detected population and the precise marker-expressions of the cells therein all demonstrate how FLARE could be used for high-speed automated identification of rare populations in cytometric data.

**Fig 3.**
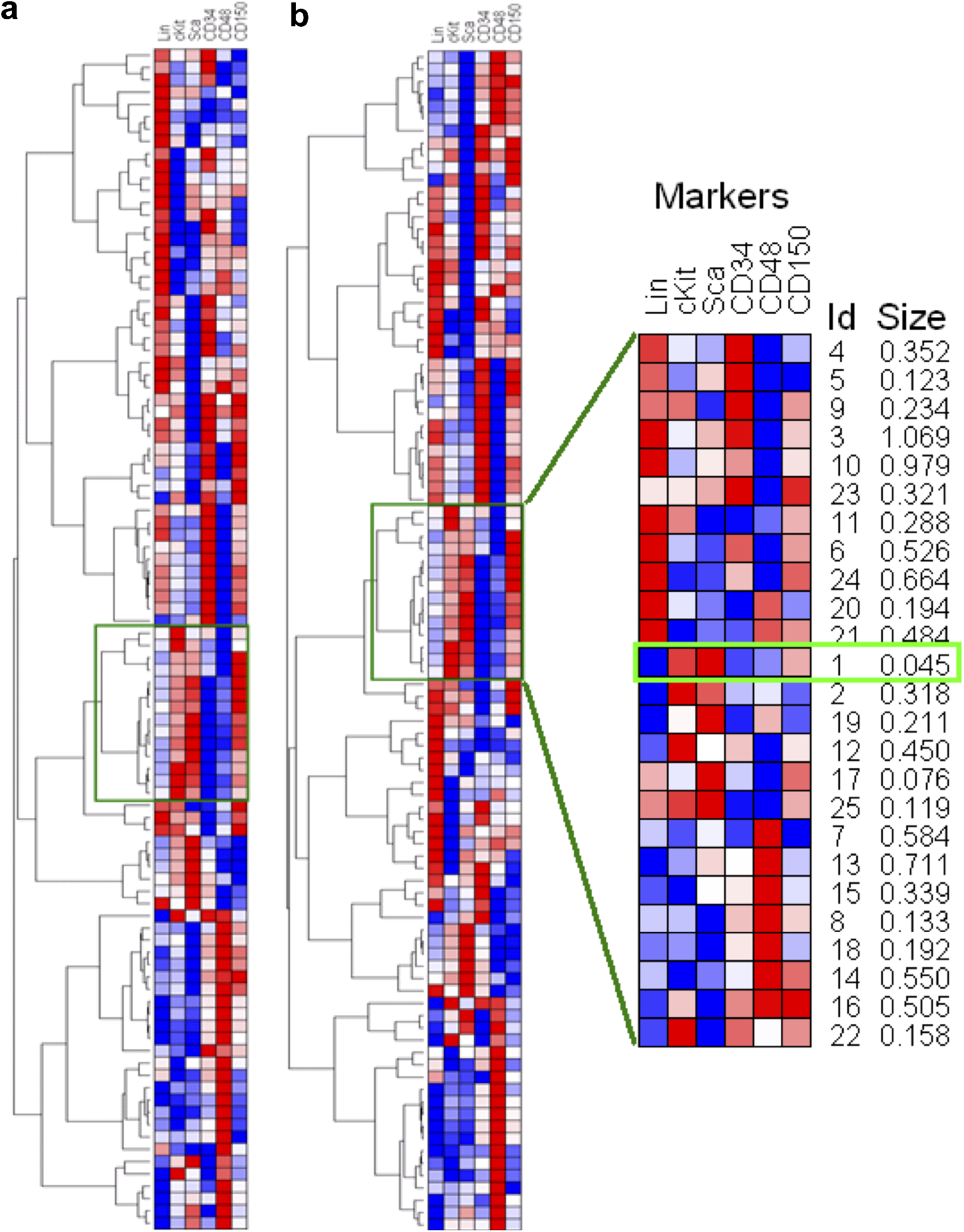
FLARE identifies rare HSC population in murine bone marrow samples undergoing normal hematopoiesis. (a) In the first step, unsupervised clustering by the hierarchical Bayesian model is used to explore the different subsets in 6-dimensional marker space. In 12 blood samples (3 biological replicate mice, 4 technical replicates per specimen), FLARE identified 96 populations matched across all samples. Plots (a) and (b) show the heatmaps for two representative samples, where each row represents a population’s mean intensities for the 6 markers represented by the columns. Red/blue is used to depict high/low intensities. Thus a common region of interest, shown in green rectangle, was identified. In the 2nd step, in a new and much larger sample, FLARE zoomed into the specified region to detect the clusters therein. The uncovered hierarchy of populations is shown with a heatmap in plot (zoomed in right panel). We identified one particular population (denoted by cluster #1; light green rectangle) that has the size (0.045%) and marker-signature (Lineage^−^c-kit^+^Sca^+^ CD34^-^CD48^-^CD150^hi^) consistent with the LT-HSC cells.

Identifying a rare disease signature as a second application of FLARE during disease progression. Therefore, we utilized a model of myeloid leukemia that harbors the oncogenic fusion of the PML gene and the retinoic acid receptor alpha (RARα). These mice succumb to a lethal acute promeylocytic leukemia (APL) that can be subsequently transplanted with increasing aggressiveness [10]. We have previously characterized the cell surface phenotype which drives the APL and it closely resembles the normal promyelocyte population [11, 12]. Notably, the mature granulocytes, which differentiate from the leukemic stem cell population (LSC) are unable to transplant the disease. The manual gating strategy for this population is challenging since these cells express low levels of lineage markers and are sequentially gated for CD34/c-Kit and then Gr1/ FcgRIIb (Lin^lo^ c-kit^+^ CD34^hi^ Gr^mid^ FcgRIIb^+^). This population in a normal bone marrow is approximately 1% of the live bone marrow cells and increases to 5-6% in the leukemic mice (Fig 4a).

**Fig 4.**
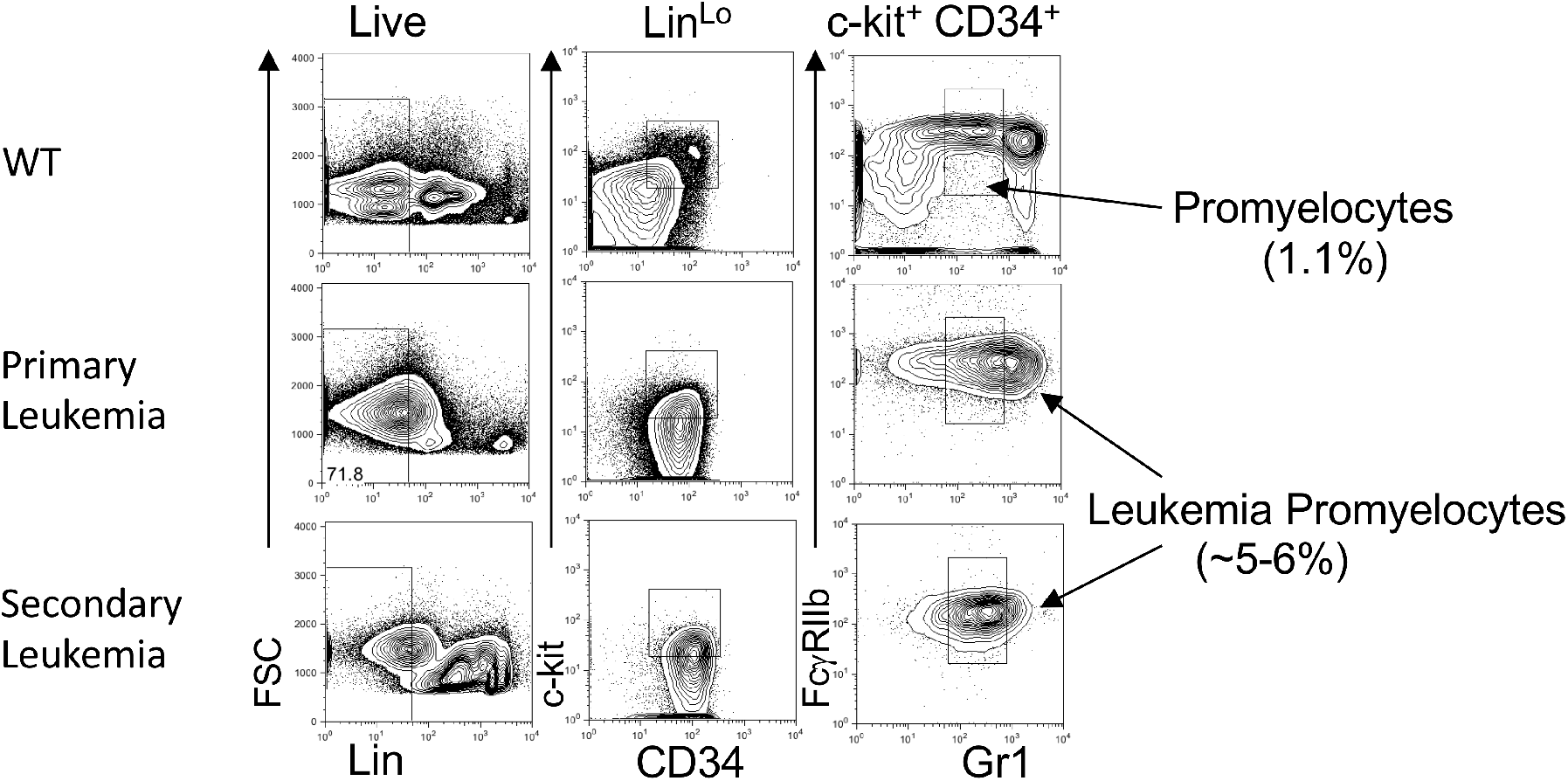
FLARE in PML-RAR*α* transgenic mouse model. Wildtype (C57BL6) bone marrow, primary leukemic mice with PML-RAR*α* transgenic and secondarily transplanted mice were analyzed by flow cytometry (and gated as previously described [11]). Forward scatter (FSC), Lineage staining (Lin) are gated serially from left panels to right. Experimentally determined surface phenotype of leukemic promyleocytes are gated (right panel), and frequency of this population is shown among live cells.

By running FLARE on live-gated cells stained with markers for lineage, c-kit, CD34, Gr-1 and FcgRIIb we could identify 99 unique clusters that contained cells (Fig 5a). To determine if the found clusters contained the LSC population, we focused on the populations with greater than 1-fold change in their proportions compared to the normal mice. This allowed us to “zoom” into 44 clusters of hematopoietic cells that increased during leukemia progression which were visualized with a heatmap (Fig. 5b). We identified a block of clusters that matched with the known surface phenotype of the promyelocytes and the LSCs, which we named the “Promyelocyte Signature” (Clusters 99, 57, 39, 25, 50 in the green box, Fig. 3c). We found the promyelocyte cluster to represent 1% of the live cells in the normal mice, which increased to 6-8% in the leukemic mice (Fig 5b). FLARE successfully identified this rare population and demonstrates the utility for tracking changes within phenotypically defined populations during disease progression. Using parallel computing, this was accomplished in under 5 minutes.

**Fig 5.**
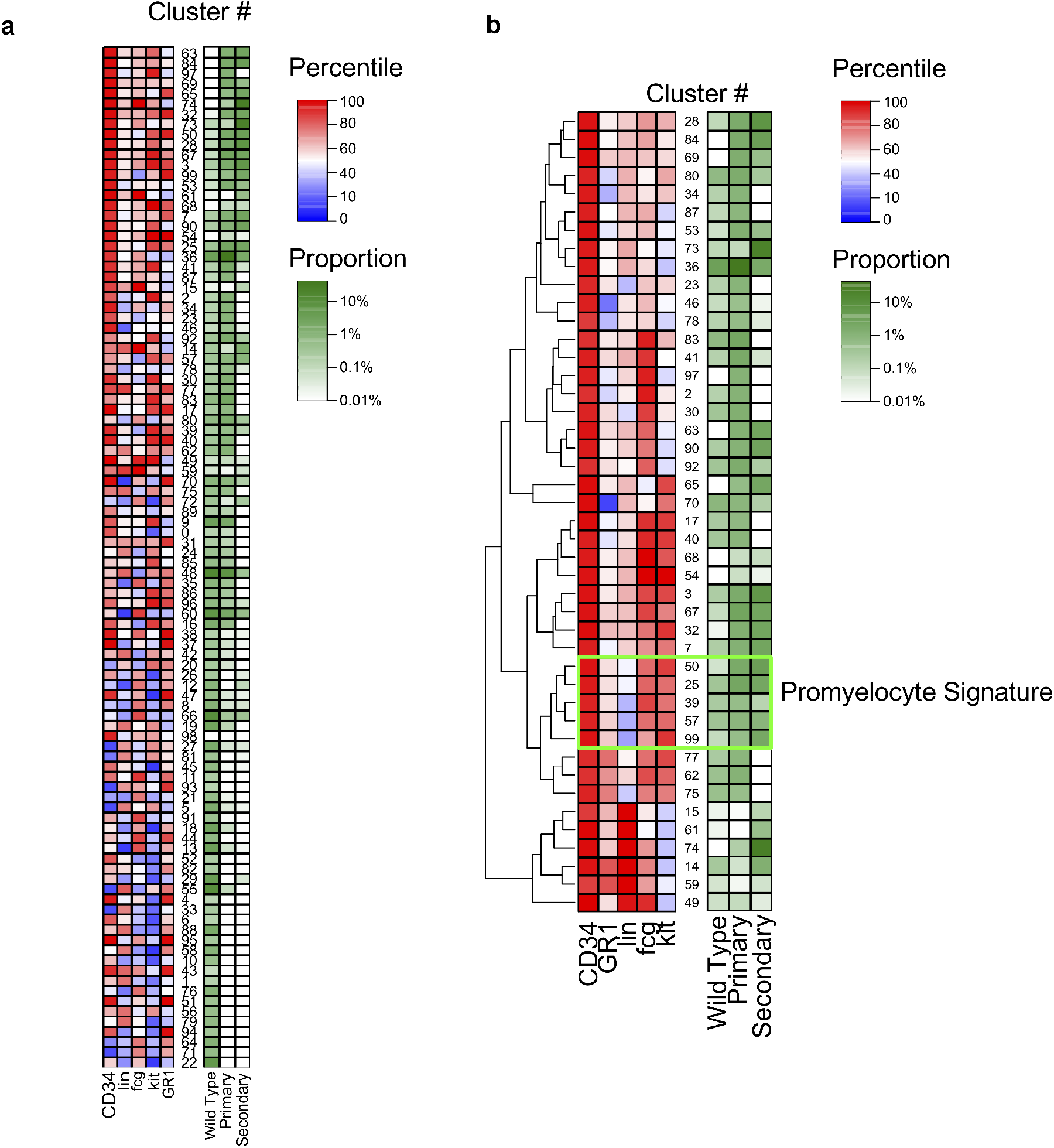
FLARE in PML-RARα transgenic mouse model. Flow cytometric data from mice in panel (a) were analyzed with FLARE and a heat map of all 99 clusters with a surface phenotype and the proportion of the particular clusters among the live cells. The populations with greater than 1-fold change in their proportions from primary or secondary leukemia compared to wild type bone marrow cells are shown in panel (b). Green box indicates clusters (50, 25, 39, 57 and 99) that contain the previously experimentally derived known surface phenotype of the leukemic promyelocytes (Lin^lo^CD34^hi^c-Kit^hi^Gr1^mid^FcgRIIb^hi^). The left panel represents the percentile of each flow staining parameter ranked among the 99 clusters. The right panel indicates proportion of live-gated cells within each cluster.

FLARE offers several distinct advantages specifically for characterization of rare populations. First, FLARE shares information across multiple samples in a hierarchical Bayesian model (Fig 1) to identify cell populations in all samples or in only part of samples. Unlike common clustering methods, FLARE does not need a priori specification of the optimal number of clusters in data, which gives it an advantage while searching samples which may contain populations ranging from significantly big to extremely rare. Instead, FLARE automatically allows an initial mixture model with a large number of components to become sparse as the inconsistent clusters are removed and the actual number of clusters used to fit the data is learned in the process. In practice, FLARE can be viewed as an efficient approximation to Dirichlet Process Mixture (DPM) models, which have been used in the past [13]. Of course, FLARE uses the strategy of information sharing for fitting robust models by verifying the rare clusters across samples. Simulation results showed that FLARE achieves favorable estimation performance over alternative methods (Supplementary Fig S2-S5).

Further, FLARE is fit with a Variational Bayes (VB) approach (see Section A of the Supplementary) which provides computationally efficient and accurate estimation of all the latent variables, i.e., the output of the model. Furthermore, FLARE modeling is parallelized with careful consideration on workload balancing in a distributed computing environment (Supplementary Algorithm 1 and Supplementary Fig S1). It achieves almost linear speedup given more computational nodes (Supplementary Fig S6), making it truly scalable for large datasets.

Identification of rare cell subsets - while establishing their correspondence across multiple samples - can (a) reveal, in an unsupervised way, the overall structure among the populations, both big and small, with respect to each other in every sample, and thereby (b) provide contextual information that helps in supervised dissection of the chosen regions of interest in the marker-space to characterize the rare populations with further precision. Such progressive “zooming in” capability of FLARE mimics the strength of sequential manual gating. An important advantage of FLARE’s Bayesian design is that it can be made to systematically zoom into interesting regions or populations by a priori specifications. Thus FLARE can perform increasingly finer clustering using the same mathematical model, which can again match and verify the finer subpopulations across multiple samples. This allows FLARE to combine the benefits of an unsupervised clustering method with supervised analysis of manual sequential gating. We illustrated these aspects of FLARE using a multi-step analysis of a hierarchy of cell populations as observed in 2 datasets based on (i) normal hematopoiesis in mice, and (ii) oncogenic progression in a mouse model. Further examples of FLARE analyses of secondary (Treg) and simulated datasets along with the performance results are described in the Supplementary Materials.

## Conclusion

In summary, the hierarchical design and distributed variational estimation allows FLARE to share information about corresponding clusters across samples, and quickly detect a variety of populations, including considerably rare ones, in an unsupervised manner. In the process, it efficiently searches the high-dimensional marker-space to reveal the underlying population structure. Thereupon it can progressively concentrate its search within regions of interest and also perform supervised analysis of subpopulations similar in principle to manual gating except FLARE does it in high-dimensions and with mathematical rigor. In our future work we look forward to embedding this step into FACS systems for real-time sorting of the desired cells. Since the multi-parametric population signatures reported by FLARE are quantitative and precise, however rare the underlying events may be, it helps to verify and eventually standardize definitions of specific cellular species, allow objective extraction, and facilitate reproducible cytometric analysis. Finally, the parallel estimation algorithm in FLARE is currently implemented using Message Passing Interface (MPI) and can be readily adapted to popular distributed computing platforms.

## Supporting information

**S1 Fig. Topography of data storage among cluster nodes.** Each rectangle represents a cluster node, with each column consisting of nodes from a particular sample. The edges of this graph connect nodes which must communicate with each other. The nodes of each column form a fully connected subgraph to show the communication done within each sample. Similarly, each column has representative node that participates in summations over all samples. The dotted edges represent the fact that based on the data, there can be an arbitrary number of samples and nodes per sample.

**S2 Fig. The Adjusted Rand Index of each method on the synthetic datasets.** We use the hard clustering results of the subject who has the small clusters to compute the ARIs against true clustering assignment.

**S3 Fig. Visualization of clustering results in synthetic data.**

**S4 Fig. Maximum Jaccard index on Treg dataset.**

**S5 Fig. Maximum detection accuracy on Treg dataset.**

**S6 Fig. Speedup rate and load balancing efficiency.** The top panel shows the speedup rate of our parallel inference algorithm using increasingly more cluster nodes. The bottom panel shows the load balancing efficiency. The balancing efficiency is calculated using equation (S32). With more nodes, the data are more evenly distributed so that the balancing efficiency keeps increasing.

**S7 Fig. Maximum detection accuracy on Treg dataset.**

**File. Supplemental Materials for ‘High-Speed Automatic Characterization of Rare Events in Flow Cytometric Data’.** Further details on Variational Inference, Parallel Inference, and Experimental Results.

