## Supplementary Materials for "High-Speed Automatic Characterization of Rare Events in Flow Cytometric Data"

### Supplemental Materials for 'High-Speed Automatic Characterization of Rare Events in Flow Cytometric Data' (Y. Qi, et al.)

#### A Variational Inference

In this section, we present a variational approach to efficiently learn this model from data.

Since there is no analytical solution to compute, the exact posterior distributions of the latent variables, a fast compilation solution is necessary. Given the large-volume of a typical cytometric data and the high-dimensionality nature of multiparametric measurements, classical Monte Carlo methods, such as Markov Chain Monte Carlo (MCMC), can be computationally costly. Thus, we resort to fast approximate Bayesian inference; in particular, we apply the Variational Bayesian (VB) method to calculate the approximate posterior distributions.

The idea of the variational inference method is to use a simpler distribution to approximate the exact posterior distribution. Specifically, the log marginal distribution can be decomposed as

$$\ln p(\mathbf{X}) = \mathcal{L}(q) + KL(q, p) \tag{S1}$$

It consists of two parts: the lower bound $\mathcal{L}(q)$ and the KL divergence between $p$ and $q$:

$$\mathcal{L}(q) = \int q(\mathbf{Z}) \ln \left\{ \frac{p(\mathbf{X}, \mathbf{Z})}{q(\mathbf{Z})} \right\} \mathrm{d}\mathbf{Z} \tag{S2}$$

$$KL(q, p) = -\int q(\mathbf{Z}) \ln \left\{ \frac{p(\mathbf{Z}|\mathbf{X})}{q(\mathbf{Z})} \right\} \mathrm{d}\mathbf{Z} \tag{S3}$$

where $X$ represents the data and $Z$ represents the random variables for which we need to calculate the posterior.

The distribution $q(\mathbf{Z})$ is the approximation of the true posterior $p(\mathbf{Z}|\mathbf{X})$. To this end, some assumptions should be exerted to the approximate posterior $q(\mathbf{Z})$. One commonly used assumption is that $q$ can be factorized as:

$$q(\mathbf{Z}) = \prod_{i=1}^{L} q_i(\mathbf{Z}_i) \tag{S4}$$

Hence, by minimizing (S3) given (S4), we can obtain the optimized form of each factor of $q(\mathbf{Z}_i)$ by:

$$\ln q_j(\mathbf{Z}_j) = \mathbb{E}_{i \neq j}[\ln p(\mathbf{X}, \mathbf{Z})] + \text{const} \tag{S5}$$

In the case of our model, we assume $q$ can be factorized as

$$q(\boldsymbol{\mu}, \boldsymbol{\Lambda}, \boldsymbol{\eta}, \boldsymbol{\Omega}, \mathbf{h}, \boldsymbol{\pi}) = q(\boldsymbol{\mu})q(\boldsymbol{\eta})q(\boldsymbol{\Lambda})q(\boldsymbol{\Omega})q(\mathbf{h})q(\boldsymbol{\pi}) \tag{S6}$$

Then by using (S5), we can obtain the optimized approximated posteriors as follows:

$$q(\mathbf{h}^{(m)}) = \prod_{n=1}^{N_m} \prod_{k=1}^{K} (r_{nk}^{(m)})^{h_{nk}^{(m)}} \tag{S7}$$

$$q(\boldsymbol{\pi}^{(m)}) = C(\boldsymbol{\alpha}^{(m)}) \prod_{k=1}^{K} (\pi_k^{(m)})^{\alpha_k^{(m)}-1} \tag{S8}$$

$$q(\boldsymbol{\mu}_k^{(m)}) = \mathcal{N}(\boldsymbol{\omega}_k^{(m)}, \boldsymbol{\Gamma}_k^{(m)}) \tag{S9}$$

$$q(\boldsymbol{\eta}_k) = \mathcal{N}(\boldsymbol{\xi}_k, \boldsymbol{\Upsilon}_k) \tag{S10}$$

$$q(\boldsymbol{\Lambda}_k^{(m)}) = \mathcal{W}(\boldsymbol{\Lambda}_k^{(m)}|\boldsymbol{\Psi}_k^{(m)}, \sigma_{m:k}) \tag{S11}$$

$$q(\boldsymbol{\Omega}_k) = \mathcal{W}^{-1}(\boldsymbol{\Omega}_k|\boldsymbol{\Phi}_k, \boldsymbol{\nu}_k) \tag{S12}$$

The parameters in these distributions are:

$$r_{nk}^{(m)} = \frac{\rho_{nk}^{(m)}}{\sum_{j=1}^{K} \rho_{nj}^{(m)}} \tag{S13}$$

$$\ln \rho_{nk}^{(m)} = (\frac{1}{2}\mathbb{E}[\ln |\boldsymbol{\Lambda}_k^{(m)}|] - \frac{1}{2}\mathbb{E}[(\mathbf{x}_n^{(m)} - \boldsymbol{\mu}_k^{(m)})^T \boldsymbol{\Lambda}_k^{(m)} (x_n^{(m)} - \boldsymbol{\mu}_k^{(m)})] - \frac{D}{2}\ln 2\pi) + \mathbb{E}[\ln \pi_{m:k}] \tag{S14}$$

$$\alpha_k^{(m)} = \alpha_0 + N_k^{(m)} \tag{S15}$$

$$N_k^{(m)} = \sum_{n=1}^{N_m} r_{nk}^{(m)} \tag{S16}$$

$$\boldsymbol{\omega}_k^{(m)} = (N_k^{(m)}\mathbb{E}[\boldsymbol{\Lambda}_k^{(m)}] + \beta_0 \mathbf{I})^{-1}(N_k^{(m)}\mathbb{E}[\boldsymbol{\Lambda}_k^{(m)}]\mathbb{E}[\mathbf{x}_k^{(m)}] + \beta_0 \mathbb{E}[\boldsymbol{\eta}_k]) \tag{S17}$$

$$\Gamma_k^{(m)} = (N_k^{(m)}\mathbb{E}[\boldsymbol{\Lambda}_k^{(m)}] + \beta_0 \mathbf{I})^{-1} \tag{S18}$$

$$\boldsymbol{\xi}_k = (\epsilon_0 \boldsymbol{\xi}_0 + \beta_0 \sum_{m=1}^{M} \mathbb{E}[\boldsymbol{\mu}_k^{(m)}])/(\epsilon_0 + \beta_0 M) \tag{S19}$$

$$\boldsymbol{\Upsilon}_k = ((\epsilon_0 + \beta_0 M)\mathbf{I})^{-1} \tag{S20}$$

$$\begin{aligned}(\boldsymbol{\Psi}_k^{(m)})^{-1} =& \sigma_0 \mathbb{E}[\boldsymbol{\Omega}_k^{-1}] + N_k^{(m)}(\boldsymbol{S}_k^{(m)} + \\ & (\mathbb{E}[\mathbf{x}_k^{(m)}] - \mathbb{E}[\boldsymbol{\mu}_k^{(m)}])(\mathbb{E}[\mathbf{x}_k^{(m)}] - \mathbb{E}[\boldsymbol{\mu}_k^{(m)}])^T)\end{aligned} \tag{S21}$$

$$\boldsymbol{S}_k^{(m)} = (\sum_{n=1}^{N_m} r_{nk}^{(m)}(\mathbf{x}_k^{(m)} - \mathbb{E}[\mathbf{x}_k^{(m)}])(\mathbf{x}_n^{(m)} - \mathbb{E}[\mathbf{x}_k^{(m)}])^T)/N_k^{(m)} \tag{S22}$$

$$\sigma_k^{(m)} = N_k^{(m)} + \sigma_0 \tag{S23}$$

$$\boldsymbol{\Phi}_k = \boldsymbol{\Phi}_0 + \sigma_0 \sum_{m=1}^{M} \mathbb{E}[\boldsymbol{\Lambda}_k^{(m)}] \tag{S24}$$

$$\nu_k = \nu_0 + M\sigma_0 \tag{S25}$$

And the expectations in the above equations are:

$$\mathbb{E}[\mathbf{x}_k^{(m)}] = \sum_{n=1}^{N_m} r_{nk}^{(m)} \mathbf{x}_n^{(m)} / N_k^{(m)} \tag{S26}$$

$$\mathbb{E}[\ln |\boldsymbol{\Lambda}_k^{(m)}|] = \sum_{i=1}^{D} \boldsymbol{\psi}(\frac{\sigma_k^{(m)} + 1 - i}{2}) + D \ln 2 + \ln |\boldsymbol{\Psi}_k^{(m)}| \tag{S27}$$

$$\begin{aligned}&\mathbb{E}[(\mathbf{x}_n^{(m)} - \boldsymbol{\mu}_k^{(m)})^T \boldsymbol{\Lambda}_k^{(m)} (\mathbf{x}_n^{(m)} - \boldsymbol{\mu}_k^{(m)})] \\ =& D/\beta_0 + \sigma_k^{(m)}(\mathbf{x}_n^{(m)} - \boldsymbol{\omega}_k^{(m)})^T \boldsymbol{\Psi}_k^{(m)} (\mathbf{x}_n^{(m)} - \boldsymbol{\omega}_k^{(m)}) + tr(\boldsymbol{\Gamma}_k^{(m)} \boldsymbol{\Psi}_k^{(m)})\end{aligned} \tag{S28}$$

$$\mathbb{E}[\ln \pi_k^{(m)}] = \psi(\alpha_k^{(m)}) - \psi(\sum_{j=1}^{K} \alpha_j^{(m)}) \tag{S29}$$

$$\mathbb{E}[\boldsymbol{\Lambda}_k^{(m)}] = \sigma_k^{(m)} \boldsymbol{\Psi}_k^{(m)} \tag{S30}$$

$$\mathbb{E}[\boldsymbol{\eta}_k] = \boldsymbol{\xi}_k, \;\; \mathbb{E}[\boldsymbol{\mu}_k^{(m)}] = \boldsymbol{\omega}_k^{(m)}, \;\; \mathbb{E}[\boldsymbol{\Omega}_k^{-1}] = \boldsymbol{\Phi}_k^{-1} \nu_k \tag{S31}$$

where $\psi(\cdot)$ is the *digamma* function—the logarithmic derivative of the gamma function $\psi(x) = \frac{d}{dx} \ln \Gamma(x) = \Gamma'(x)/\Gamma(x)$.

### B Parallel Inference

Thousands of cells are processed by a flow cytometer every second, which results in an extremely high volume of data to analyze. For example, one of our data sets contains nearly 2.6 million cells spread across 14 samples. Due to the complexity of our model and the size of flow cytometry data, there is a critical need to develop a parallel algorithm which can take advantage of the processing power in a large-scale computer cluster. While the sequential version of FLARE was implemented using Matlab, the parallel version is implemented using C++ and MPI.

A computer cluster consists of many separate nodes, i.e. computers, connected via a fast local area network. Additionally, each node may contain a multi-core processor. This allows us to devise a two-level parallelization scheme to analyze the data. At the first level of parallelization, we divide the data amongst the cluster nodes. Hence, each cluster node is responsible for a portion of the raw data as well as maintaining any parameters associated with that data. For example, all sample means ($\boldsymbol{\omega}$) and sample precision matrices ($\boldsymbol{\Psi}$) for sample 1 need to be stored on any cluster node which contains data from sample 1. In an effort to minimize repeated storage, we impose the restriction that each node must only store data from a single sample. Additionally, every cluster node may have a multi-core processor, which enables us to implement a second level of parallelization based on the number Gaussian components ($K$). Many of the parameters we infer are indexed by $k$, e.g. the prototype mean $\boldsymbol{\xi}$ is really a set of $K$ prototype means (one mean for each component). Therefore, we can use the set of processor cores to optimize the variational inference parameters for each value of $k$ in parallel.

#### B.1 Data Partition

Each iteration of the parallel inference algorithm alternates several times between computation and communication phases. All nodes must complete their current computation phase before the next one one can begin. Therefore, the total execution time is dependent on the node with the highest computational load (the node that take the longest). The goal of load balancing is to minimize the largest computational load of the nodes in the computer cluster.

Suppose we have a computer cluster consisting of a total of $W$ nodes. We must now find some way to distribute the data among these $W$ nodes that minimizes the total execution time. A naive approach would be to evenly divide the $N$ data points so that each node is responsible for $N/W$ data points. However, the volume of data assigned to each node is not the only factor that influences computation time. A node must also maintain all parameters associated with its data. For instance, a node with data from samples 1 and 2 will need to maintain means and precision matrices for both of these samples, whereas a node that only has data from sample 1 will maintain means and precision matrices for sample 1 only. Therefore the time spent spent optimizing distribution parameters can be reduced by restricting each node to data from a single sample.

**Algorithm 1** Balance the computation load across the available cluster nodes

```
function NodeBalance
    if The number of nodes is less than the number of samples then
        Error!
    end if
    Assign one node to each sample.
    while There are unassigned nodes do
        Assign a node to the sample with the highest load.
    end while
end function
```

We can think of the balancing problem in this way: we have $W$ nodes available, and a subset of these nodes ($W_1$) must be assigned to sample 1, another subset ($W_2$) must be assigned to sample 2, and so on for all $M$ samples. The data of a particular sample is divided evenly among the nodes assigned that sample. The load for each node assigned to a particular sample is equal to $N_m/W_m$. Algorithm 1 gives us a greedy strategy to minimize the load on the sample with the largest load. In order for this algorithm to function correctly, we must declare a larger number of nodes than there are samples.

We define the data partition efficiency by

$$\text{balance} = \frac{\text{load}_{\text{opt}}}{\text{load}_{\text{max}}} \tag{S32}$$

Since the optimum balance would have the same computational load on each node, we define $\text{load}_{\text{opt}}$ by

$$\text{load}_{\text{opt}} = \lceil W/N \rceil \tag{S33}$$

Where

$$N = \sum_{m=1}^{M} N_m \tag{S34}$$

Also, we define the maximum load ($\text{load}_{\text{max}}$) by

$$\text{load}_{\text{max}} = \underset{m=1,..,M}{\text{argmax}}\,(W_m) \tag{S35}$$

We know that $\text{load}_{\text{max}} \leq \text{load}_{\text{opt}}$ because any time the load differs from the optimum, we must have some node with a larger load than $\text{load}_{\text{opt}}$, and some other node with load smaller than $\text{load}_{\text{opt}}$. Therefore, an optimally balanced set of nodes will give us a data partition efficiency score of 1, and any non-optimally balanced set of nodes will give us a score less than 1. Also since the true computational cost is dependent on the slowest node, we use the node with the largest load to define the data partition efficiency.

#### B.2 Organization of Parameters Across Cluster Nodes

The raw data is not the only information we must store. The variational inference method gives us a set of parameters we must iteratively optimize. Namely, these parameters are $\mathbf{r}$, $\boldsymbol{\rho}$, $\alpha$, $N$, $\boldsymbol{\omega}$, $E[\mathbf{x}]$, $\boldsymbol{\Gamma}$, $\boldsymbol{\xi}$, $\boldsymbol{\Psi}$, $\boldsymbol{S}$, $\sigma$, $\boldsymbol{\Phi}$, and $\nu$. We can divide these parameters into 3 separate categories based of how they are indexed. The first category includes all parameters indexed by sample and by data point. These parameters include $r$, $\rho$, and the raw data $x$. The second category includes all parameters indexed by sample. These parameters include $\alpha$, $N$, $\boldsymbol{\omega}$, $E[\mathbf{x}]$, $\boldsymbol{\Gamma}$, $\boldsymbol{\Psi}$, and $S$. The third category includes all parameters which are not indexed by sample or by data point. These parameters include $\boldsymbol{\xi}$, $\boldsymbol{\Phi}$, and $\nu$. To show how these three groups of parameters are stored on the cluster, we define three new parameters, $A$, $B$, and $C$.

$A$ is used to represent the first category and is indexed in the following way. $A$'s superscript is indexed by sample, so $A^{(m)}$ includes $r^{(m)}$, $\rho^{(m)}$, and $\mathbf{x}^{(m)}$ for all $m = 1, ..., M$. Furthermore, each $A^{(m)}$ is split up into $W_m$ different parts, where $W_m$ is the number of nodes assigned to sample $m$. Each of these $W_m$ parts contains an equal portion of the $N_m$ data points in sample $m$. So, $A_1^{(m)}$ includes $r_1^{(m)}$ to $r_{d_m}^{(m)}$, $\rho_1^{(m)}$ to $\rho_{d_m}^{(m)}$, and $\mathbf{x}_1^{(m)}$ to $\mathbf{x}_{d_m}^{(m)}$, where $d_m = N_m/W_m$. Similarly $A_2^{(m)}$ contains the next $N_m/W_m$ indices of $r^{(m)}$, $\rho^{(m)}$, and $\mathbf{x}^{(m)}$, and so on for all $W_m$ portions.

The topographies of $B$ and $C$ are much simpler. $B$ is used to represent the second category and is indexed by sample. Hence, $B^{(m)}$ includes $\alpha^{(m)}$, $N^{(m)}$, $\boldsymbol{\omega}^{(m)}$, $E[\mathbf{x}]$, $\boldsymbol{\Gamma}^{(m)}$, $\boldsymbol{\Psi}^{(m)}$, and $S^{(m)}$ for all $m = 1, ..., M$. $C$ is used to represent the third category and is not indexed.

With the parameters $A$, $B$, and $C$ in hand, we can visualize the overall cluster topography as shown in Fig. S1. Using this topography, the calculations of equations (S13) and (S14) are split up among every node with no repeated calculation. The calculation of equations (S15), (S17), (S18), and (S23) can be done with no communication. However the calculation of these equations is repeated on every node assigned to a particular sample, e.g. the calculation of these equations for sample 1 is repeated on all nodes assigned to sample 1. Equations (S16), (S26), and (S22) all involve a summation indexed from 1 to $N_m$. The nodes of sample $m$ all calculate their partial sum using their portion of the data, then communicate to calculate the total sum. The calculations for each sample can be done simultaneously. Lastly, equations (S19) and (S24) involve a summation indexed from 1 to $M$. To perform this calculation a representative node from each sample is chosen to contribute its partial sum. Each of these representative nodes then communicate their results to the rest of the nodes assigned to their respective samples.

#### B.3 Further Parallelization using p-threads

Each cluster node may have a multi-core processor. With the exception of equation (S13), each of the parameter update equations for variational inference are indexed by Gaussian components, where each $k = 1, ..., K$ is independent. Therefore, all of these equations may be updated simultaneously using p-threads.

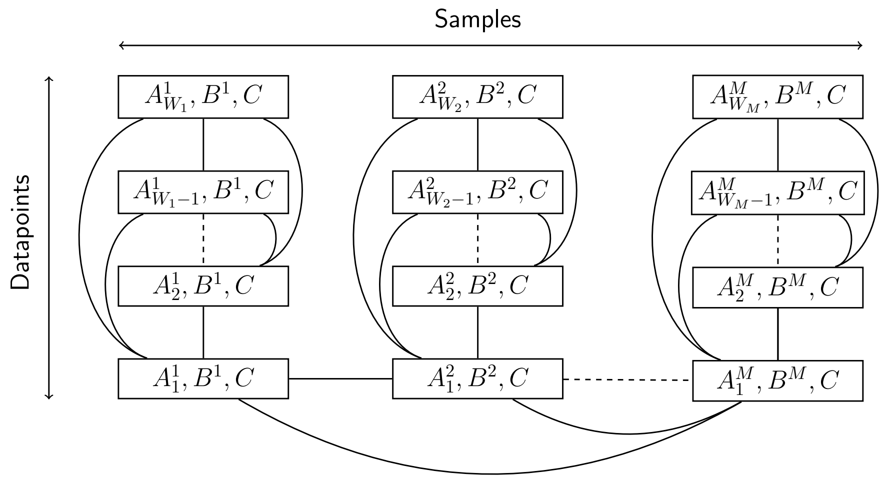

Figure S1: Topography of data storage among cluster nodes. Each rectangle represents a cluster node, with each column consisting of nodes from a particular sample. The edges of this graph connect nodes which must communicate with each other. The nodes of each column form a fully connected subgraph to show the communication done within each sample. Similarly, each column has representative node that participates in summations over all samples. The dotted edges represent the fact that based on the data, there can be an arbitrary number of samples and nodes per sample.

### C Experimental Results

#### C.1 Synthetic data

We perform the experiment on synthetic and real flow cytometry datasets.

First, using synthetic datasets where we know the ground truth, we compare our method with three state-of-the-art flow cytometry analysis methods, FlowMeans [1], FlowClust [2] and SamSpectral [3]. FlowMeans is based on the classic Kmeans algorithm; FlowClust uses a multi-variate $t$ mixture model; and SamSpectral uses spectral clustering as its core engine. We set the initial number of clusters $K$ to be the same for all methods. The three comparison algorithms use merging techniques to combine similar clusters after the clustering procedures. With a Dirichlet prior that encourages sparse mixture components, our model automatically prunes unnecessary clusters based on the data. It is worth noting that all the comparison methods do not explicitly differentiate multiple samples to find rare cell clusters. By contrast, our model can identify the common clusters across samples and differentiate rare clusters that may only present in few samples.

In our simulation, we generate three large clusters in three samples. One of the three samples has an additional rare cluster close to one of the large clusters. Each large cluster in one sample has 100 data points, and the rare cluster has only 10 points. The center and variance of the big clusters are generated according to the graphical model in Fig 1 of the main document—we generate the prototype means and covariances by sampling from equations 6 and 7 and use them to sample the means and covariances of each sample by equations 4 and 5. We vary the dimension of the data from 2 to 16 and repeat the experiment 50 times. To assess the accuracy of the clustering result, we compute the Adjusted Rand Index (ARI) [4] between the clustering results and the true clusters. The higher the ARI, the more accurate the clustering results. The average ARIs for each method are reported in Fig. S2. Our method achieved the highest ARI compared to other methods.

We visualize the clustering results of our method and the competing methods with a 2D synthetic dataset in Fig. S3. In this figure, each row is a different clustering method, and each column is a sample. The sample having the rare cluster is in the last column. Different clusters are represented by different colors. Our method is the only method that differentiates the rare cluster from the three large clusters as shown in the fourth

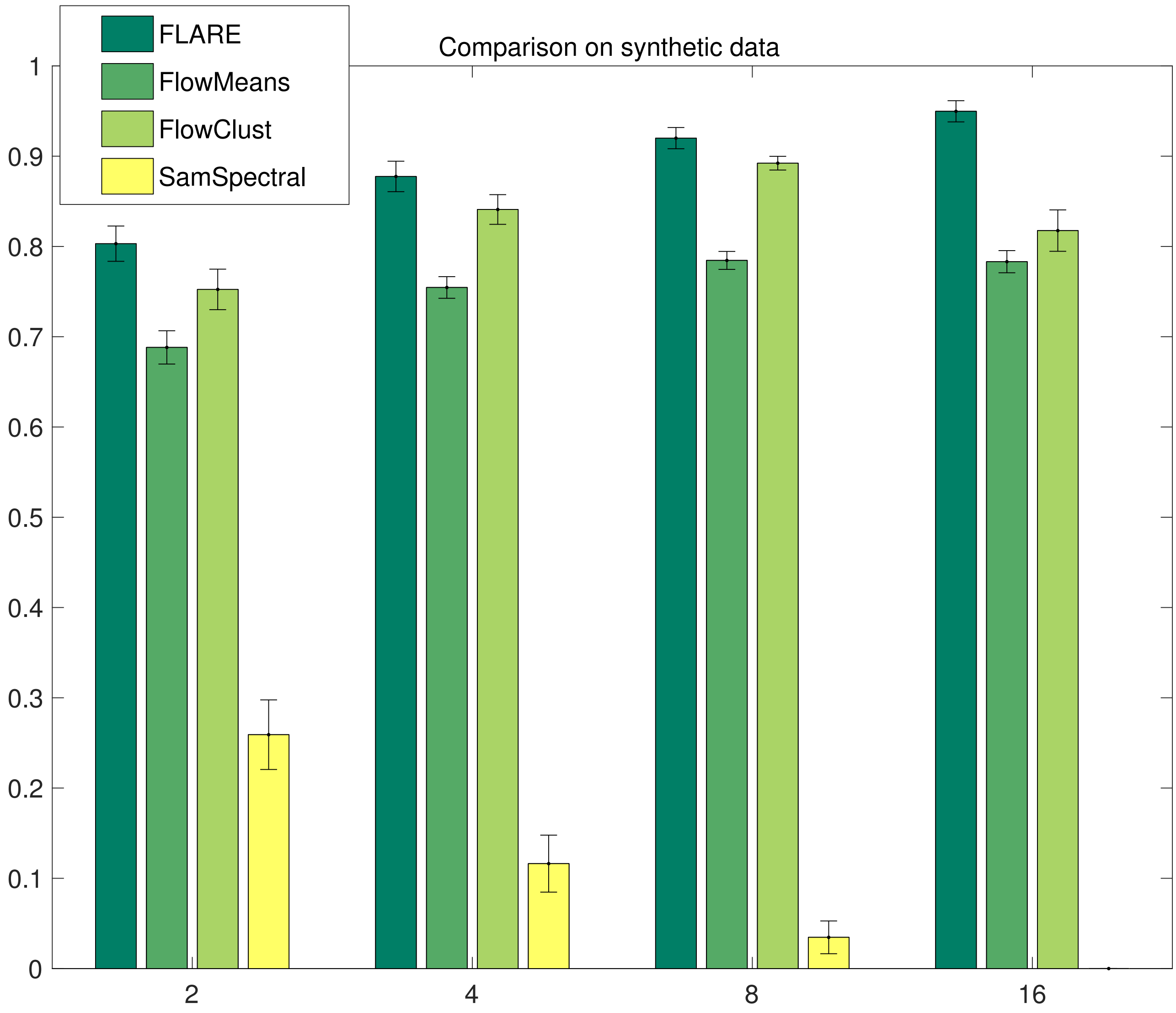

Figure S2: The Adjusted Rand Index of each method on the synthetic datasets. We use the hard clustering results of the subject who has the small clusters to compute the ARIs against true clustering assignment.

column.

#### C.2 Treg data

We also test our method on a small real-world data set, Treg. There are two samples in this dataset. Each sample has $\sim 28,000$ cells. A small artificial T-cell cluster ($\sim 300$ cells) is added to one of the samples. The task is to identify this small T-cell cluster, i.e. the target cluster. To assess the clustering methods, we compute the Jaccard index between each computed cluster and the target cluster. For a cluster $C$ and the target cluste $T$, Jaccard index is defined as

$$J(C,T) = \frac{|C \cap T|}{|C \cup T|}. \tag{S36}$$

For each method, we report the maximum Jaccard index computed between the target cluster and the all computed clusters. The result is shown in Fig. S4, with standard error over 10 runs with different initializations. Again, our method outperforms the alternative methods significantly.

The denominator of Jaccard index includes the number of data points in the computed clusters. We are also interested in the proportion of the data points in the target cluster that fall in the computed cluster. To this end, we define *detection accuracy* as the following quantity

$$D(C,T) = \frac{|C \cap T|}{|T|}. \tag{S37}$$

and report the maximum $D(C,T)$ for all the computed clusters $C$ for each method in Fig. S5. FLARE consistently outperforms the other three methods.

#### C.3 Speedup Results

A big advantage of FLARE is that we are able to analyze millions of cells with our parallel inference algorithm on computer clusters within minutes. We tested this parallel inference algorithm using a cluster called Carter, which is maintained by Purdue's Rosen Center for Advanced computing. Carter consists of a total of 660 nodes, Each of which contains two 8-core Intel Xeon-E5 processors. The nodes each have 56 Gbps FDR Infiniband connections. To facilitate communication between nodes we use the Message Passing Interface (MPI) version 1.4.4. As shown in Fig. S6, we are able to achieve nearly linear speedup: just under 5 seconds for the oncogenic signature data set, 265 seconds for the scaled hematopoiesis data.

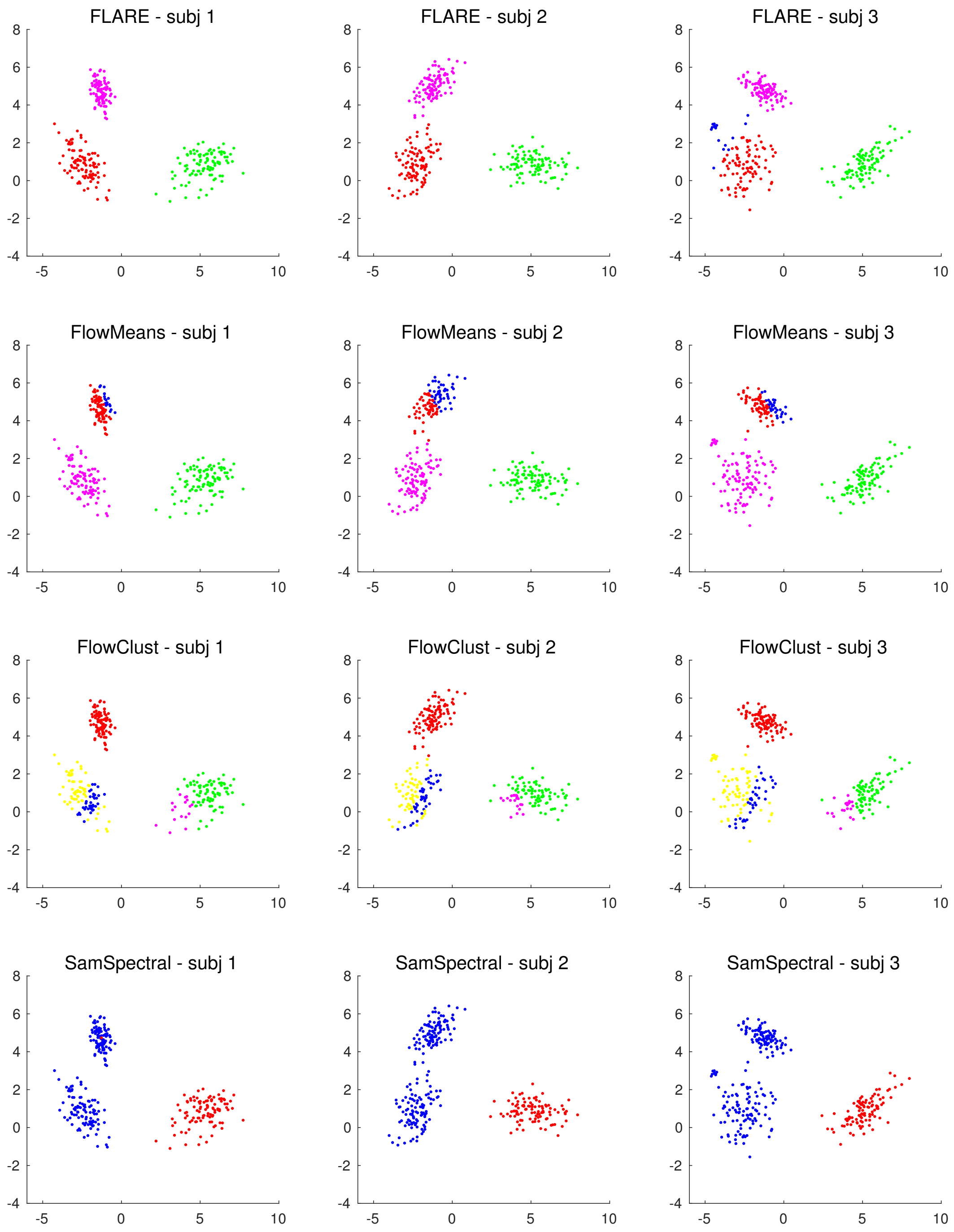

Figure S3: Visualization of clustering results in synthetic data.

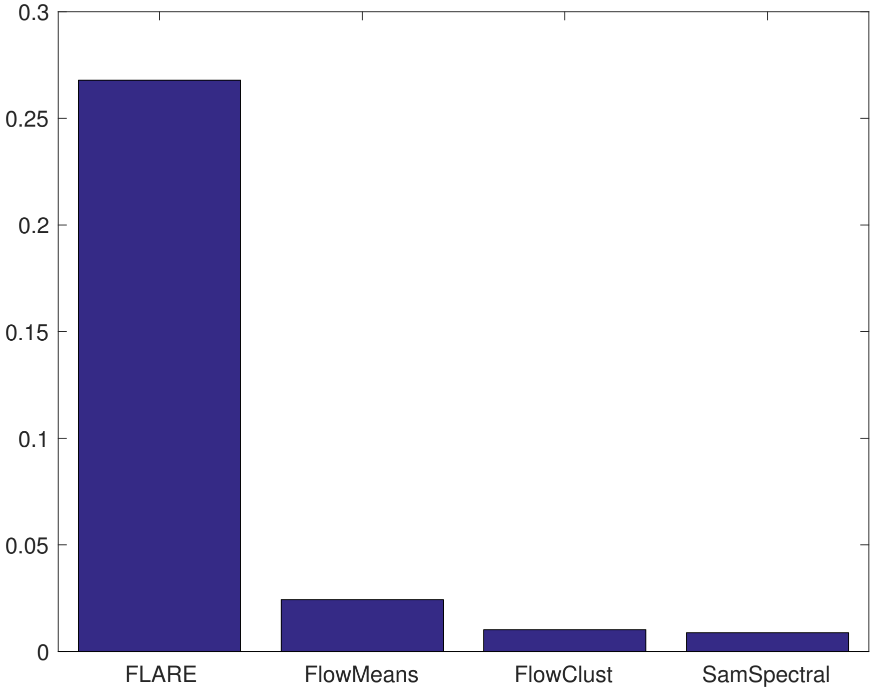

Figure S4: Maximum Jaccard index on Treg dataset.

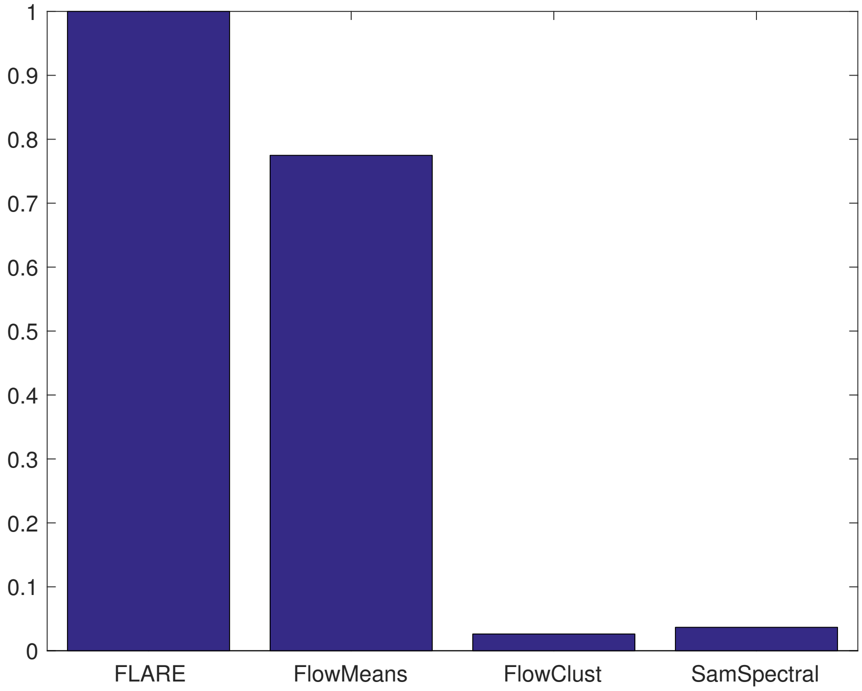

Figure S5: Maximum detection accuracy on Treg dataset.

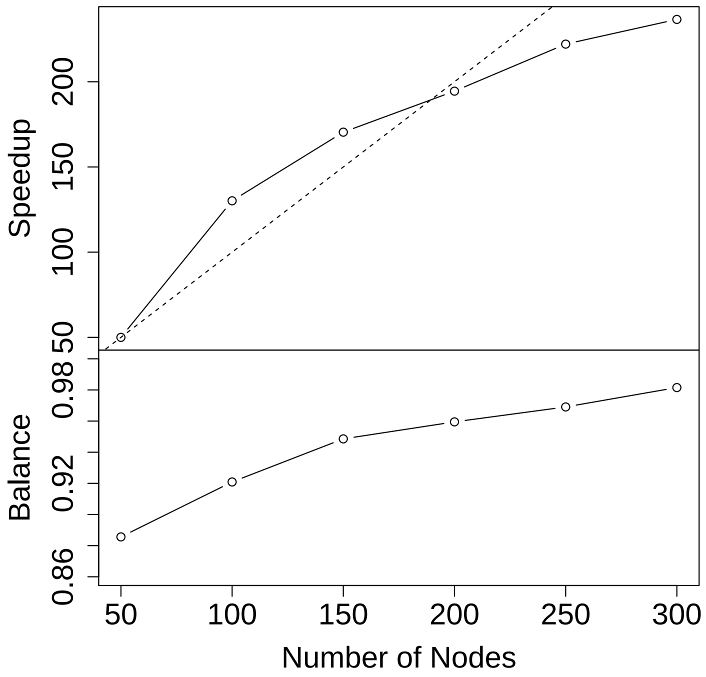

Figure S6: The top panel shows the speedup rate of our parallel inference algorithm using increasingly more cluster nodes. The bottom panel shows the load balancing efficiency. The balancing efficiency is calculated using equation (S32). With more nodes, the data are more evenly distributed so that the balancing efficiency keeps increasing.
